# Schizophrenia-Linked 3q29 Deletion Produces Hyperdopaminergic Signaling in the Dorsal Striatum

**DOI:** 10.64898/2026.11.13.766445

**Authors:** Sindhu Sriramoji-Virdi, Rebecca M Pollak, Anna Tischer, Michael Michaelides, Jennifer Gladys Mulle, Miriam E. Bocarsly

**Affiliations:** Department of Pharmacology, Physiology and Neuroscience, New Jersey Medical School, Rutgers University, Newark, New Jersey, USA; Center for Advanced Biotechnology and Medicine, Robert Wood Johnson Medical School, Rutgers University, Piscataway, New Jersey, USA; National Institute on Drug Abuse, Intramural Research Program, NIH, Baltimore, Maryland, USA; Department of Psychiatry, Robert Wood Johnson Medical School, Rutgers University, Piscataway, New Jersey, USA; Rutgers Brain Health Institute, Rutgers University, Piscataway, New Jersey, USA

## Abstract

Dopamine dysregulation has long been recognized as a central feature of schizophrenia (SZ), yet the mechanisms linking genetic risk to dopaminergic dysfunction remain poorly understood. While pharmacological models have historically been used to induce hyperdopaminergic states and recapitulate select clinical features of SZ, these approaches do not model the underlying disease biology. In contrast, genetically informed models enable investigation of dopaminergic abnormalities as emergent consequences of bona fide disease-associated mechanisms. Here, we examined a mouse model of the 3q29 deletion. In humans, this 1.6 Mb deletion is typically *de novo* and confers a greater than 40-fold increased risk for SZ, representing the single largest known molecular risk factor for the disorder. Using this model, we identified a robust hyperdopaminergic phenotype characterized by elevated evoked dopamine release selectively within the dorsal striatum. Mechanistically, this phenotype appears to arise from a PKC-mediated reduction in dopamine transporter (DAT) availability, impairing dopamine clearance and prolonging dopaminergic signaling. Together, these results establish a direct link between a high-risk SZ mutation and striatal dopamine dysfunction, while also highlighting PKC-DAT signaling as a potential therapeutic target for treatment-resistant neuropsychiatric symptoms associated with 3q29 deletion.

## Introduction

Dopamine dysregulation has been recognized as a central feature of schizophrenia (SZ) for over 75 years^1–3^. Historically, pharmacological models have been used to recapitulate specific clinical features of SZ through reverse translation, such as inducing hyperdopaminergic states^4,5^. And although valuable for probing symptom-related processes, these approaches do not model the underlying disease biology and instead reproduce isolated phenotypic features. That is, these models can answer questions related to the consequences of hyperdopaminergia but cannot address the causal mechanisms by which this state arises in the first place.

In contrast, the current study leverages a mouse model harboring a SZ-associated genetic deletion, enabling the examination of dopamine dysfunction as an emergent consequence of bona fide genetic risks and directly linking dopaminergic abnormalities to disease etiology without experimentally imposing circuit perturbations. Here, we focus on a 1.6 mega-base deletion at chromosome 3 (3q29 deletion; hg19, chr3:195725000-197350000), a typically *de novo* mutation in humans, which is associated with a range of neuropsychiatric phenotypes including intellectual disability, autism, anxiety, and most notably a >40-fold increase in the risk of developing SZ^6–11^. As the single-largest molecular risk factor for SZ^3^, the 3q29 deletion is a powerful tool for interrogating the unknown neurobiology underlying SZ-associated pathology.

In the current study, we utilized a mouse model of the 3q29 deletion^12^. Designed on a C57B/6N background, the mouse was generated using CRISPR/Cas9-mediated excision of the syntenic region on mouse chromosome 16^12^. As expected, there was ∼50% reduction in expression of nearly all the genes contained within the deletion segment, as well as many of the protein products, validating the utility of the system to explore underlying molecular and cellular mechanisms^12^.

In the current study we find that the 3q29 deletion mouse model exhibits spontaneous elevated evoked dopamine release, selectively in the dorsal striatum. We further find that this phenotype appears to be driven by a PKC-mediated reduction in dopamine transporter (DAT) availability. DAT is a membrane protein that regulates dopaminergic signaling by clearing dopamine from the synaptic cleft back into presynaptic neurons, thereby controlling the strength and duration of the signal. While there is a literature suggesting DAT dysfunction may be a driving factor in the hyperdopaminergic state of SZ^13^, this is the first study to show increased dopaminergic function in the SZ-associated 3q29 deletion mouse model and provide a novel mechanism with possible treatment potential.

## Methods

### Mice

All procedures were approved by the Animal Care and Use Committees of Rutgers University and conducted according to specifications of the National Institutes of Health as outlined in the Guide for the Care and Use of Laboratory Animals. Male and female mice were group-housed on a 12 h light-dark cycle (lights on at 7:00 AM) and given *ad libitum* access to water and standard rodent chow. Mice (3q29del) harboring a heterozygous deletion of the region syntenic to human 3q29 on chromosome 16 of the C57BL/6N background were generated and genotyped as previously described^12^. Littermate mice lacking the heterozygous deletion were used as controls. Experimentation began at 6 weeks of age. The number of animals per group used in experiments ranged between 5 and 25 and is indicated in figure legends.

### Fast-Scan Cyclic Voltammetry

#### Brain slice preparation

Mice were deeply anesthetized with isoflurane, decapitated, and brains were rapidly harvested and placed in warm cutting solution (in mM): 225 sucrose, 13.9 NaCl, 26.2 NaHCO_3_, 1 NaH_2_PO_4_, 1.25 glucose, 2.5 KCl, 0.1 CaCl_2_, 4.9 MgCl_2_, and 3 kynurenic acid. Brains were sliced sagittaly using a vibratome (240 μm) in warm cutting solution. Brain sections containing dorsomedial striatum or the nucleus accumbens were allowed to recover in oxygenated ACSF containing (in mM): 124 NaCl, 2.5 KCl, 2.5 CaCl_2_, 1.3 MgCl_2_, 26.2 NaHCO_3_, 1 NaH_2_PO_4_, and 20 glucose (∼310-315 mOsm) at 33⁰C for 30-60 min and maintained at room temperature until recording.

#### Fast-scan cyclic voltammetry recordings

Fast-scanning cyclic voltammetry was performed in the dorsomedial striatum and the nucleus accumbens shell from 3q29del and control mice to determine levels of evoked dopamine. Carbon fibers (7μm diameter) were inserted into glass pipettes to create carbon fiber electrodes (CFEs), which were hand-cut to expose ∼150μm of the fiber past the pipette tip. Before use, CFEs were backfilled with 3M KCl internal solution. CFEs were conditioned before recordings with a triangular voltage ramp (-0.4 to 1.2 and back to -0.4 V versus Ag/AgCl reference at 400 V/s) delivered every 15ms. During recordings, CFEs were held at -0.4 V versus Ag/AgCl while the same triangular voltage ramp was delivered every 100ms. Dopamine transients were evoked by a single pulse (0.2 ms) of electrical stimulation administered every 2 minutes by a glass micropipette filled with ACSF that was placed near the CFE tip. To generate input-output curves, the intensity of electrical stimulations was varied (10, 20, 40, 60, 80, 100, 150, 200, 300, or 400 µA) with each episode. Experiments were conducted in ACSF and maintained at 31-33℃. In a subset of experiments, slices were treated with Clorgiline (MAO-A inhibitor, 100nM), Selegiline (MAO-B inhibitor, 100 nM), Tolcapone (COMT inhibitor, 100nM), Disulfiram (ALDH1A1 inhibitor, 100 µM), GBR12909 (DAT blocker, 5 µM), Tetrabenazine (VMAT2 inhibitor, 100 µM), or LY333531 (PKCß inhibitor, 100-200 µM). Data were acquired and analyzed using a Chem-Clamp amplifier (Dagan) and I/O board (National Instruments) with a custom-written software VIGOR using Igor Pro (Wavemetrics) running mafPC (courtesy of M.A. Xu-Friedman). All data were analyzed with VIGOR and peak current amplitude of dopamine was converted to concentrations according to the post-experimental calibration.

### DAT mRNA Measurements

Quantitative polymerase chain reaction was used to measure relative mRNA levels in striatal brain tissue. Mice were anesthetized with isoflurane and decapitated. Brains were removed, and the striatum was dissected on ice using a 1 mm coronal matrix, placed in RNAlater, homogenized, and total RNA was purified using RNeasy Plus Mini kit (QIAGEN). cDNA was synthesized using iScript Reverse Transcription Supermix (Biorad). Slc6a3 (Mm00438388_m1) and InsR (Mm00439688_m1) TaqMan Gene Expression Assays (Applied Biosystems) were used to determine relative mRNA expression compared to the endogenous control gene viniculin (Mm00447745_m1). Samples were run in triplicate and in parallel with negative controls using the StepOnePlus Real-Time PCR system (Applied Biosystems). The cycling conditions were: initial hold at 95°C (20 s), 40 cycles of 95°C (1 s) and 60°C (20 s). Relative mRNA levels were calculated using the ΔΔCt method.

### Autoradiography

DAT receptor density was measured in the striatum of mice using autoradiography. Whole brains were flash-frozen in 2-methylbutane for later sectioning (20 μm) on a Cryostat (Leica) and thaw mounted onto ethanol-washed glass slides. Slides were preincubated for 20 min in ice-cold washing buffer (50 mM Tris-HCl, pH 7.4 with 120 mM NaCl), then incubated in the same buffer conditions for 120 min containing [^3^H]WIN 35428 (10 nM, 82.5 Ci/mmol, Revvity). Another set of slides containing consecutive sections was incubated in the same conditions in the presence of cocaine HCl (100 μM, Tocris) to determine nonspecific binding. Slides were washed 2 x 1 min in ice-cold Tris-HCl (50 mM) and rinsed for 30 sec in ice-cold deionized water. Then the slides were dried overnight, placed in a hypercassette (Amersham Biosciences), and covered with a BAS-TR2025 Storage Phosphor Screen (Fujifilm). A slide containing Carbon-14 Standards (American Radiolabeled Chemicals Inc.) was exposed simultaneously for quantitative radiometric analysis. The slides were exposed to the screen for 12 days and imaged using a Typhoon biomolecular imager (Cytiva). Images were calibrated and analyzed using ImageJ 1.51j8 (NIH). Regions of interest (ROI; 8 ROI per region and animal) were drawn freehand based on neuroanatomical landmarks. The non-specific binding of each ROI was subtracted from total binding to calculate specific binding (nCi/g), which was normalized to the wild-type control animals to calculate % specific binding.

### Western blot analysis

Mice were euthanized and dorsal striatum samples were collected and flash frozen. Tissue samples were lysed in RIPA buffer with protease inhibitor (Thermo Scientific 89901), homogenized and then sonicated. Protein was evaluated using a Pierce BCA protein assay (Thermofisher 23225) at a 1:5 dilution. Protein (20 µg) was boiled in loading buffer (Bio-rad #1610374; 95°C, 7 min). Samples were loaded onto TGX stain-free mini gels (Bio-Rad #4561035) and run at 100 V for 2 h. Gels were transferred to a 0.2 µm nitrocellulose membrane using the Bio-Rad Trans-Blot Turbo at 2.5 amps, 25 V for 6 min. Poncu staining (Thermo Scientific A40000279) was utilized to confirm equal protein loading across wells. Membranes were blocked in 5% BSA in tris-buffered saline solution with tween (TBST) for one hour at room temperature with rapid shaking.

Primary antibodies (Anti-Tyrosine Hydroxylase: Millipore Sigma AB152; Anti-phospho-Tyrosine Hydroxylase: Millipore Sigma AB152; Anti-Dopamine Transporter: Proteintech 22524-1-AP; Anti-phospho-Dopamine Transporter: ABCAM ab183486; Anti-Mao-A (10539-1-AP); Anti-Mao-B (12602-1-AP); and Anti-Vinculin (Cell Signaling, 1390) were diluted at 1:1000 in TBST and incubated at 4°C with gentle shaking. Anti-COMT (14754-1-AP) and Anti-ALDH1A (15910-1-AP) were diluted at 1:300 in TBST and likewise incubated at 4°C with gentle shaking. Following primary incubation, membranes were rinsed with TBST 4×10 min with rapid shaking and then incubated with secondary HRP (Gt, Thermo Fisher, 18-4818-82 or Rb, Thermo Fisher, NA9340V, 1:5000) in 5% BSA in TBST for 1 h at room temperature with gentle shaking, followed by 2×15 min TBST rinses with rapid shaking. For development, membranes were covered in ECL (GE Healthcare, 17030484) for 2 min and imaged using chemiluminescence on the Chemi-doc imaging system. For blots in which multiple proteins could be detected from a single membrane, membranes were stripped between hybridizations with a stripping buffer (10% SDS, Tris HCl, pH 6.5, and 0.8% beta-mercaptoethanol) at room temperature for 15 min. Membranes were moved to TBST and rinsed 2×15 min. Membranes were re-imaged to ensure adequate stripping before re-blocking. Primary and secondary incubations were repeated as described. Band intensity was quantified using the Image Lab software (Bio-Rad).

### Locomotor testing

Two separate cohorts of mice were used to assess locomotor activity in response to pharmacological agents. Locomotor activity was assessed during the light phase. Mice were moved to the locomotor testing room and allowed to acclimate for 30 minutes. They were then individually placed in polycarbonate chambers (20 cm H × 17 cm W × 28 cm D) equipped with infrared photobeam detectors (Columbus Instruments, Columbus, OH). Beam breaks were recorded for 1 h while mice habituated chambers, allowing for the novelty-induced dopamine response to attenuate. Mice were then administered drug (i.p.) and locomotion was measured for an additional hour. On days 1-3, all mice received saline (10 ml/kg). Cohort 1 mice were treated with the DAT inhibitor GBR-12909 (1 mg/kg, i.p.). Cohort 2 mice were treated with amphetamine (either 1 or 7.5 mg/kg, i.p. with 2 days between doses).

### Intrastriatal injection of PKCß inhibitor

Each adult mouse was placed into a stereotaxic frame under isoflurane anesthesia, and a 26-gauge bilateral guide cannula (Protech International) was implanted above the dorsal striatum (AP: +1.00, ML: ± 0.5, DV: −2.0, with relation to bregma). Cannulas were fixed in place using 2 micro-screws and dental cement (Cooralite Dental Manufacturing). A dummy stylet was placed into the guide and protected with a dust cap when not in use. Mice recovered for at least one week after surgery before experimental procedures began. On the experimental days, mice were moved to the locomotor testing room and allowed to acclimate for 30 minutes. They were then individually placed in polycarbonate chambers (20 cm H × 17 cm W × 28 cm D) equipped with infrared photobeam detectors (Columbus Instruments, Columbus, OH). Beam breaks were recorded for 1 h while mice habituated chambers, allowing for the novelty-induced dopamine response to attenuate. On days 1-3, mice were given intrastriatal injections of saline (5µl, 2.5 µl/min) and on day 4 mice received the PKCß inhibitor LY333531 (200 nM, 5 µl, 2.5 µl/min). The injector was left in place for 1 min following injection to ensure full diffusion, and mice were placed back in the boxes and locomotion was measured for an additional hour.

### Statistical analysis

Analyses were performed in Prism 10 (GraphPad). Data making three or more comparisons were analyzed using two-way ANOVA, with the addition of repeated-measures (RM) or a mixed-effects model, as appropriate (ex. voltammetry input/output curves and time courses, locomotor behavior, etc.). For RM-ANOVA, sphericity was assessed with Mauchly’s test. Significant main effects or interactions were followed up with pairwise tests corrected for multiple comparisons (Sidak or Tukey). Linear mixed effects models were utilized when multiple tissue slices from a single animal were examined. Data comparing two groups (ex. mRNA levels, protein levels, etc. between genotypes) were compared with an unpaired Student’s t-test. One sample t-test was used to determine deviations from the mean or baseline. Results were considered significant at an alpha ≤ 0.05. All data are presented as mean ± SEM and individual animal data is shown whenever possible. Full statistics are presented in the figure legends. Sample sizes were chosen based on previous research employing similar approaches and are sufficient for detecting effects while using the minimal number of animals. Data that is presented as “% of control” were normalized to the average of the control group.

## Results

### 3q29 deletion mice show elevated dopamine in the dorsal striatum

In humans, the *de novo* 3q29 deletion is associated with a >40-fold increase in the risk of developing schizophrenia (SZ)^1^. In the following studies, we used a 3q29 deletion mouse model generated on a C57BL/6N background, harboring a heterozygous CRISPR/Cas9-mediated excision of the region syntenic to human 3q29 on mouse chromosome 16 **(Figure 1A)**^12^. Because dopamine dysfunction has long been implicated in the pathophysiology of SZ^14^, we first employed ex vivo fast-scan cyclic voltammetry (FSCV), an electrochemical technique, to assess alterations in dopamine release in 3q29 deletion mice relative to littermate controls. 3q29 deletion mice showed increased levels of evoked dopamine across a range of stimulation intensities in the dorsal striatum compared to littermate controls (**Figure 1B**). The mean dopamine transients have a peak amplitude of 744.7 ± 51.0 nM in control mice versus 924.4 ± 62.8 nM for 3q29 deletion mice (p = 0.01; **Figure 1C-E**). Further, the decay of the dopamine signals was significantly different between genotypes, as indicated by differences in the calculated decay constant, Tau (0.37 ± 0.03 vs. 0.49 ± 0.04, p = 0.03; **Figure 1F**). Interestingly, the same findings did not reach statistical significance in the nucleus accumbens, suggesting regional changes in dopamine release capacity (**Figure 1G-I**). Despite no differences in overall dopamine release in the nucleus accumbens, tau was found to be elevated (p = 0.04; **Figure 1J**). No differences in dopamine release or tau were seen in the dorsal striatum or nucleus accumbens between male and female mice (**Supplemental Figure 1A-F**), supporting collapsing data across sexes for the remainder of this paper.

**Figure 1.**
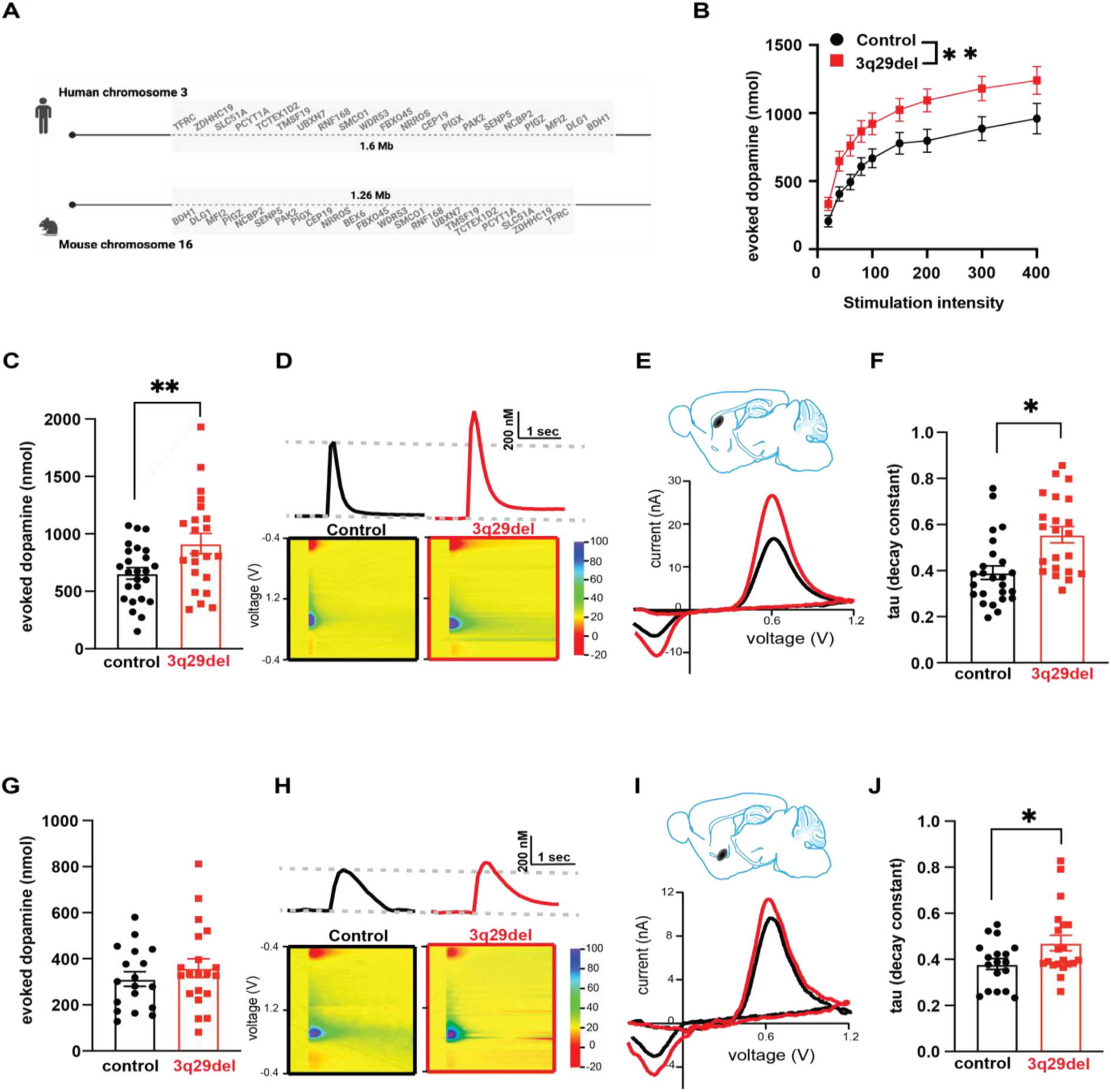
3q29 deletion mice show elevated evoked dopamine in the dorsal striatum but not the nucleus accumbens. **A.** The 3q29 region is present on human chromosome 3 [deletion coordinates (GRCh38.p12)-Chr3:195,998,129–197,623,129]. The deletion is simulated on mouse chromosome 16 [deletion coordinates (GRCm38.p3)-Chr16:31,369,117–32,634,414]. The genes within this region are presented here. **B.** Mean peak amplitude of evoked dopamine at increasing voltage stimulation intensities in the dorsal striatum of 3q29del mice (red) compared to control mice (black). Control n = 50 slices/20 mice. 3q29del n = 40 slices/18 mice. Mixed effects model: main effect of genotype F (1,88) = 7.64, p = 0.007. Main effect of stimulation intensity F(8,691) = 123.0 p<0.0001) **C.** Amplitudes of evoked dopamine signals at 200μA stimulation in the dorsal striatum for control mice (black) and 3q29del mice (red). Control n = 77 slices/25 mice. 3q29del n = 72 slices/23 mice. Mixed effects model: main effect of genotype F(1,45)=7.10, p = 0.01, no effect of slice F(6,94)=0.45, p=0.84. **D.** (Top) Representative traces of dopamine signals in the dorsal striatum of control mice (black) and 3q29del mice (red). (Bottom) Representative voltammograms evoked by a single 200μA electrical stimulation in the dorsal striatum. **E.** (Top) Shaded area represents site of recordings. (Bottom) Representative current voltage plots of 3q29del mice (red) compared to control mice (black) of the electrochemical current (background-subtracted) evoked by stimulation in the dorsal striatum showing the characteristic current peaks at 0.6 V and −0.2 V that match the oxidation and reduction potentials for dopamine. **F.** Decay constant (*tau)* of 3q29del mice (red) compared to control mice (black) in dorsal striatum. Mixed effects model: main effect of genotype F(1,45)=5.06, p = 0.03. No effect of slice F(6,94)=1.39, p=0.23. **G.** Amplitudes of evoked dopamine signals at 200μA stimulation in ventral striatum of control mice (black) and 3q29del (red). Control n = 18 slices/18 mice. 3q29del n = 20 slices/20 mice. Unpaired t-test: no effect of genotype t(36)=0.91, p = 0.37. **H.** (Top) Representative traces of dopamine signals in the ventral striatum of 3q29del mice (red) compared to control mice (black). (Bottom) Representative voltammograms evoked by a single 200μA electrical stimulation in the nucleus accumbens. **I.** (Top) Shaded area represents site of recordings. (Bottom) Representative current voltage plots of 3q29del mice (red) compared to control mice (black). **J.** Decay constant (*tau)* of 3q29del mice (red) compared to control mice (black) in dorsal striatum. Unpaired t-test: t(36)=2.16, p=0.04. Data is presented as mean ± SEM, with individual points representing data from a single mouse.

### Dopamine synthesis and degradation enzymes are unchanged in 3q29 deletion mice

We next aimed to determine mechanistic changes in the 3q29 deletion mice that could be leading to the observation of increased evoked dopamine in the dorsal striatum. There are several factors that could contribute to the hyperdopaminergic state, including changed in synthesis, degradation, or recycling (see **Figure 2A**). We initially hypothesized that increased dopamine release could be the result of changes in tyrosine hydroxylase (TH), the rate-limiting enzyme in the conversion of tyrosine to dopamine^15^. However, no differences were seen in the levels of phosphorylated serine40-Tyrosin Hydroxylase, total TH, or the ratio of the two, in striatal tissue homogenate from 3q29 deletion and control mice (**Figure 2B**).

**Figure 2.**
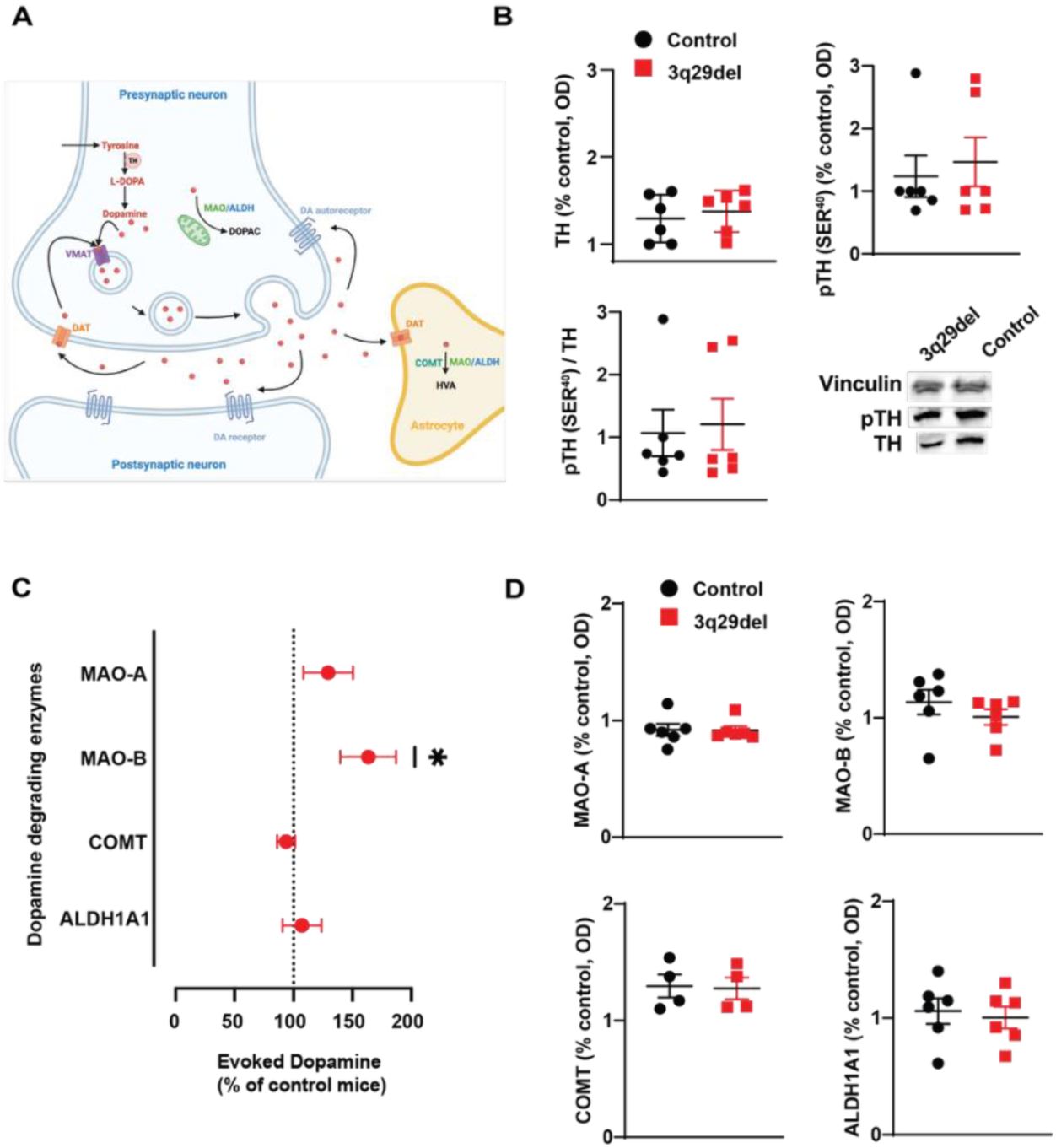
3q29 deletion mice show elevated Mao B activity in the dorsal striatum. **A.** Schematic illustrating key enzymes involved in dopamine metabolism. **B.** Protein level estimation (normalized to vinculin) of TH (unpaired t-test; t(10) = 0.56, p = 0.59), p-TH(Ser 40) (t(10) = 0.44, p =0.66) in the dorsal striatum of 3q29del mice (red) compared to control mice (black), as well as pTH/TH (t(10) = 0.26, p =0.80). Representative western blot shown. **C.** Evoked dopamine amplitudes following 200μA of voltage in tissue slices containing the dorsal striatum, following application of select enzyme inhibitors. Dashed line represents averaged values for control mice, and deviation from control is shown for 3q29del mice (red). All time courses of dopamine release in control and 3q29del mice can be found in the supplementary figures. Enzymatic washes included: 100nm Mao-A inhibitor Clorgiline (one sample *t-test* from control baseline: t(12)=1.40, p=0.19). 100nm Mao-B inhibitor Selegiline (t(12)=2.69, p=0.02). 100nm COMT inhibitor Tolcapone (t(12)=3.10, p=0.009). 100 µM ALDH inhibitor Disulfiram (t(24)=0.44, p=0.67). **D.** Protein estimation results of MAO-A (t(10)=0.07, p=0.95), Mao-B (t(10)=1.02, p=0.33), COMT (t(6)=0.16, p=0.88), and ALD (t(1 0)=0.38, p=0.71) relative to vinculin in dorsal striatum of 3q29del mice (red) compared to control mice (black) n=4-6 mice/group. Representative western blot in supplemental figures. Schematic was created using BioRender. **ALDH** Aldehyde Dehydrogenase, **COMT** Catechol-O-Methyltransferase, **DA** Dopamine, **DAT** Dopamine Transporter, **DOPAC** 3,4-Dihydroxyphenylacetic Acid, **HVA** Homovanillic Acid, **L-DOPA** Levodopa (L-3,4-dihydroxyphenylalanine), **MAO-A** Monoamine Oxidase A, **MAO-B** Monoamine Oxidase B, **pTH** Phosphorylated Tyrosine Hydroxylase, **TH** Tyrosine Hydroxylase, **VMAT** Vesicular Monoamine Transporter. Data is presented as mean ± SEM, with individual points representing data from a single mouse.

In the absence of detectable changes in TH expression, we examined the dopamine-degrading enzymes responsible for dopamine metabolism. Specifically, we focused on monoamine oxidase A (MAO-A), monoamine oxidase B (MAO-B), catechol-O-methyltransferase (COMT), and aldehyde dehydrogenase 1 family member A1 (ALDH1A1)^16^. Using FSCV, we examined changes in evoked dopamine in 3q29 deletion mice and littermate controls following the application of blockers of each of these specific dopamine degrading enzymes. In control mice, as anticipated, blocking the dopamine degrading enzymes, MAO-A, COMT, and ALDH1A1 led to a moderate increase in evoked dopamine in control mice (**Supplementary Figure 2A,C-D**). On the other hand, inhibition of MAO-B did not lead to an increase in evoked dopamine in control mice (**Supplementary Figure 2B**). In further analysis, blocking MAO-A, COMT, and ALDH1A1 in 3q29 deletion mice led to dopamine profiles similar to those seen in littermate controls (**Figure 2C**), while blocking MAO-B led to more evoked dopamine in the 3q29 deletion mice compared to littermate controls (p = 0.02). This is, perhaps, reflective of increased obligation of MAO-B in degrading the excessive dopamine levels in 3q29 deletion mice compared to littermate controls.

To better understand the role of the dopamine degrading enzymes in the 3q29 deletion mice, we looked at protein expression of the enzymes. No differences were seen in protein expression of the dopamine degrading enzymes MAO-A, MAO-B, COMT, or ALDH1A1 in striatal tissue homogenate from 3q29 deletion and control mice (**Figure 2D**). Together, these data led us to the conclusion that deficits in dopamine degrading enzymes are not driving the observed increase in evoked dopamine in the dorsal striatum of 3q29 deletion mice.

### Altered DAT function underlies increases in dorsal striatal dopamine in 3q29 deletion mice

Finding no explanation of the elevated dopamine in the function of the dopamine degrading enzymes, we turned to enzymes responsible for the reuptake and subsequent repackaging of dopamine following vesicular release (dopamine transporter (DAT) and vesicular monoamine transporter (Vmat), respectively). While no differences were seen in 3q29 deletion and littermate controls in Vmat function, which is responsible for the repackaging of dopamine back into vesicles (**Supplementary Figure 3A,B**), differences in DAT function, which is responsible for the reuptake of dopamine back into the cell, were seen (**Figure 3**). Using ex vivo FSCV, when the atypical DAT inhibitor, GBR-12909, was applied, control mice showed elevated levels of evoked dopamine in the dorsal striatum (time course seen in **Figure 3A**). However, the expected increase in evoked dopamine following GBR-12909 application in 3q29 deletion mice was highly attenuated compared to control mice (**Figure 3B**; 239 ± 47% increase from baseline in controls and 145 ± 21% increase from baseline in 3q29 deletion mice, p = 0.01). Further, in a behavioral study, where the locomotor sensitivity of the same DAT antagonist (GBR-12909) was tested in 3q29 deletion mice compared to littermate controls, we found that 3q29 deletion mice showed a trend towards locomotor suppression in response to a low dose of GBR-12909 compared to controls (**Figure 3C,D**). Together, this data suggests that DAT function is diminished in 3q29 deletion mice, compared to littermate controls. To determine whether the diminished DAT function is driven by differences in density of DAT, we examined the binding of [^3^H]WIN-35428, a DAT-selective radioligand, in 3q29 deletion mice compared to littermate controls, and found a decrease in DAT binding in the dorsal striatum (p = 0.02; **Figure 3E**).

**Figure 3.**
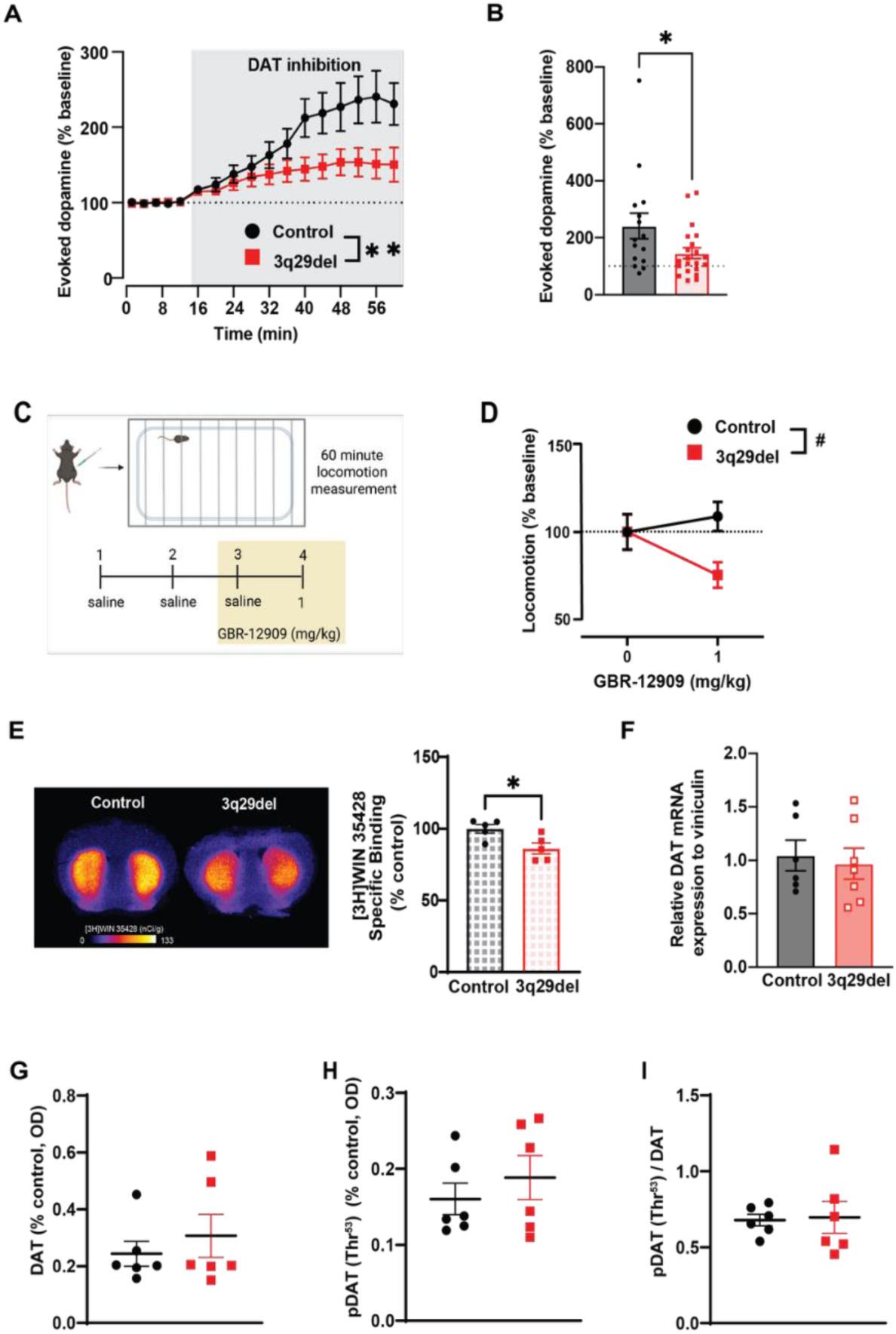
3q29deletion mice show insufficient dopamine transporter activity and decreased DAT binding relative to control mice in the dorsal striatum. **A.** Normalized time course of dopamine peak amplitudes following the DAT inhibitor, GBR-12909 (5 µM), in tissue samples containing the dorsal striatum from 3q29del and control mice. (2-way RM ANOVA: interaction effect (F(16,490)=6.32, p<0.0001, main effect of genotype F(1,36)=10.12, p=0.003, main effect of drug F(16,490) = 22.64, p<0.0001). **B.** Amplitudes of evoked dopamine signals at 200µA in dorsal striatum following incubation in GBR-12909 (5 µM) for control mice (black, 15 slices/6 mice) and 3q29del mice (red, 22 slices/11 mice; unpaired t-test t(35)=2.58, p = 0.01). **C.** Schematic showing experimental timeline where locomotion was measures following either saline or the DAT inhibitor GBR-12909 (i.p.). **D.** Mean normalized locomotor activity in 3q29del mice (red) compared to control mice (black) following saline (0 mg/kg) or GBR-12909 (1 mg/kg; 2-way RM ANOVA: effect of genotype F(1,20) = 3.54 p=0.07; post hoc pairwise comparison suggests emerging differences between genotypes (p=0.02). **E.** Representative sections depicting binding of DAT radioligand [^3^H]WIN 35428 and % specific binding in the dorsal striatum of 3q29del mice (red) compared to control mice (black; n=5 mice/group; unpaired t-test t(8)=2.82, p = 0.02). **F.** Relative DAT mRNA in the dorsal striatum of 3q29del mice (red, n=7 mice) compared to control mice (black; n=6 mice; unpaired t-test t(11)=0.37, p=0.72). **G.** DAT protein, relative to vinculin, in dorsal striatum of 3q29del mice compared to control mice (black; n=6/group; unpaired t-test t(10)=0.40, p=0.70). **H.** Levels of pDAT (Thr53) relative to vinculin in the dorsal striatum of 3q29del mice (red) compared to control mice (black; n=6/group; t(10)=0.79, p=0.45). **I.** Ratio of p-DAT (Thr53)/DAT in the dorsal striatum of 3q29del mice (red) compared to control mice (black; n=6 group; unpaired t-test t(10)=0.16 p=0.88). Representative western blot can be found in supplemental figures. Data is presented as mean ± SEM, with individual points representing data from a single mouse. Schematic was created using BioRender.

Despite reduced DAT binding in 3q29 deletion mice compared to littermate controls, DAT mRNA and protein expression were no different in the dorsal striatum across genotypes (**Figure 3F,G**). Further, no differences were seen in the levels of phosphorylated Threonine53-DAT or when it was normalized to total DAT in striatal tissue homogenate from 3q29 deletion and control mice (**Figure 3H,I**). This potentially indicates that DAT is present in its phosphorylated state but not expressed in the membrane, or reduced DAT binding could also be driven by decreased DAT affinity.

### Elevated dopamine in 3q29 deletion mice is associated with changes to PKCβ function

To determine what might be leading to the decreased binding of DAT, we explored several factors that are known to influence the membrane presence of DAT including activation of the insulin, muscarinic, and nicotinic receptors^17–19^. However, no differences were seen in the levels of these proteins or the dopaminergic release in response to blocking these receptors in the dorsal striatum between the 3q29 deletion mice and controls (**Supplementary Figure 5**).

Still searching for the mechanism underlying the decreased DAT function, and driving the hyperdopaminergic state, we turned to an early observation made in the 3q29 deletion mice. In the original 3q29 deletion mouse characterization paper, it was observed that 3q29 deletion mice showed a blunted locomotor response to amphetamine^12^, which we replicated here (p = 0.004; **Figure 4A,B**). The literature indicates that sensitivity to amphetamines can indicate the efficacy of protein kinase C-beta (PKCβ) in regulating dopamine signaling^20^. When activated, PKCβ phosphorylates and then ubiquitinates DAT, leading to its internalization from the cell membrane, ultimately decreasing surface expression, decreasing dopamine reuptake and affecting neurotransmission^21^. In wildtype mice, blocking PKCβ interferes with dopamine homeostasis, with increased membrane DAT leading to an overall decrease in dopamine available for release. We observed a decrease in evoked dopamine using FSCV following the application of the PKCβ antagonist, LY333531 (100 μM), in control mice. However, in 3q29 deletion mice, LY333531 showed no change in evoked dopamine at 100 μM. When the concentration of LY333531 was increased to 200 μM, both genotypes similarly showed depressed levels of evoked dopamine (**Figure 4C**; p = 0.03). To assess DAT function, we analyzed tau, the decay constant of dopamine transients, which is commonly used as an index of dopamine clearance kinetics^22^. In control mice, incubation with 200 μM LY333531 produced a robust increase in tau, indicating slower dopamine clearance (**Figure 4D**, p = 0.04). In contrast, 3q29 deletion mice exhibited only a modest increase in tau, suggesting impaired DAT function and a reduced sensitivity to PKCβ-mediated regulation. In a final experiment, mice were prepared with cannula aimed at the dorsal striatum for the local administration of LY333531. Following LY333531, control mice showed a suppression in locomotor activity, while 3q29 deletion mice showed no change in locomotor activity (**Figure 4E,F**; p = 0.03). All together, these data lead us to propose that PKCβ is upregulated in 3q29 deletion mice, leading to a decrease in DAT function and increased dopamine in the synapse (**Figure 4G**).

**Figure 4.**
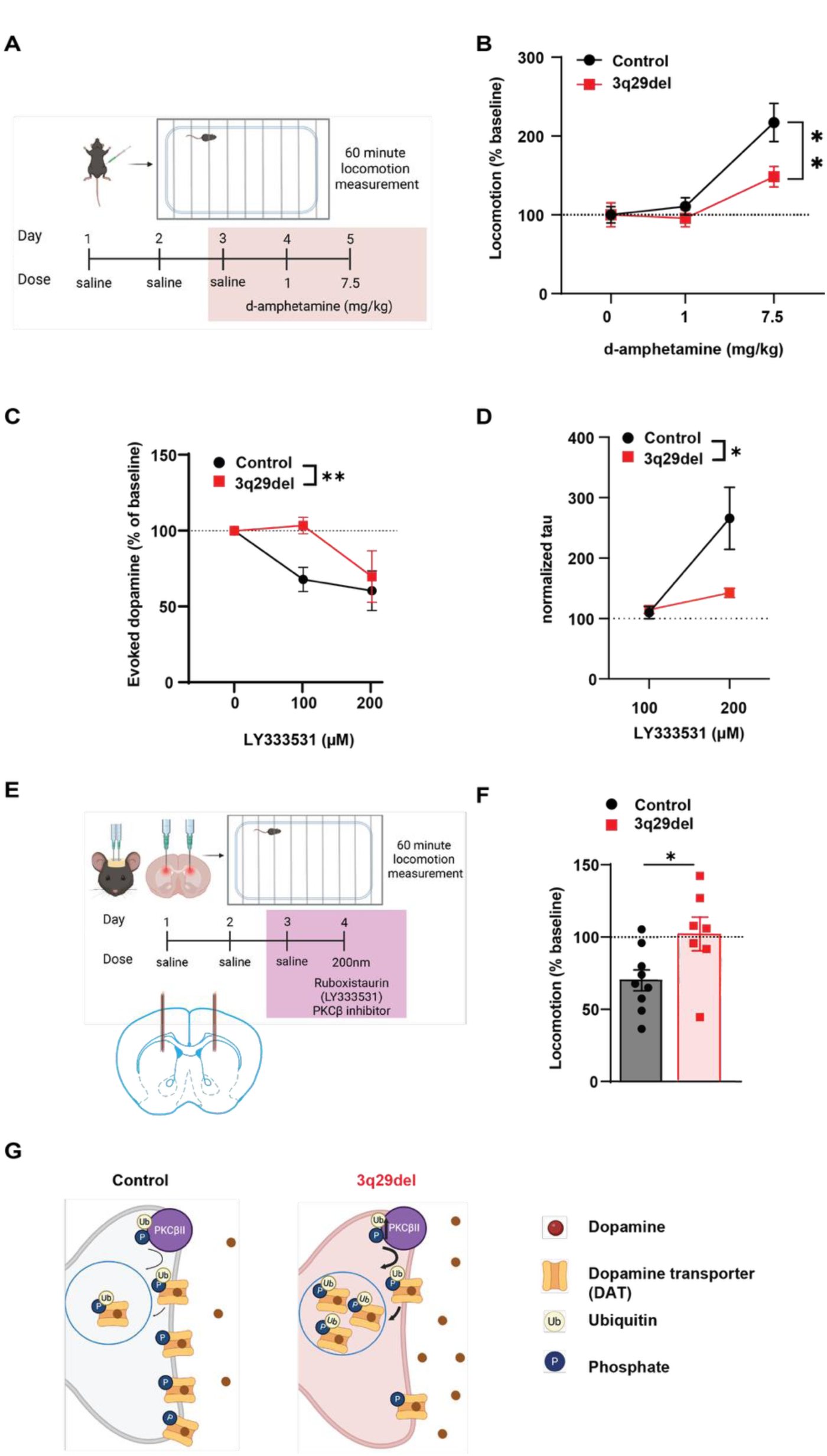
3q29 deletion mice show increased levels of PKCβ activation. **A.** Schematic showing experimental timeline where locomotion was measures following either saline or d-amphetamine administration (i.p.). **B.** Mean locomotor activity 30 minutes following d-amphetamine, normalized to saline, for 3q29del mice (red) compared to control mice (black; n=10 mice/group). 2-way RM ANOVA: interaction effect F(2,38) = 4.78, p=0.01; main effect of dose F(2,38) = 26.94, p<0.0001. Post hoc pairwise comparisons show differences between genotypes in locomotion following 7.5mg/kg of d-amphetamine (p = 0.004). **C.** Mean evoked dopamine, as a percent of baseline, following ruboxistaurin (LY333531) treatment of brain slices containing the dorsal striatum from 3q29del mice (red) compared to control mice (black, 9 slices/3 mice per group). 2-way RM ANOVA main effect of dose F(2, 40) = 8.97, p = 0.0006; main effect of genotype F(1,40) = 5.32, p= 0.03. Post hoc pairwise comparisons indicate differences between genotypes following 100μm ruboxistaurin (p=0.008). **D**. From the same experiment, we determined the normalized decay constant, tau, following ruboxistaurin incubation. Data is normalized to pre-treatment baseline. 2-way RM ANOVA main effect of genotype F(1,12) = 4.98, p = 0.04; main effect of dose F(1,12) = 11.80, p = 0.005. Post hoc pairwise comparisons indicate differences between genotypes following 200μm ruboxistaurin (p = 0.007). **E.** Schematic showing experimental timeline where locomotion was measures following either saline or ruboxistaurin (LY333531), a PKCβ inhibitor, injected directly into the dorsal striatum. As well as pictural representation of histology confirming injection placement. **F.** Locomotor activity for 60 minutes following intrastriatal ruboxistaurin administration, as a percent of locomotion following saline for 3q29del mice (red, n= 7 mice) compared to control mice (black, n=9 mice; unpaired t-test t(14)=2.43, p=0.03). **G.** Schematic illustrating hypothesized changes in PKCβ-meditated dopamine metabolism in 3q29del mouse model, compared to controls. Data is presented as mean ± SEM, with individual points representing data from a single mouse. Schematic was created using BioRender.

## Discussion

Elevated striatal dopamine signaling is a hallmark of schizophrenia pathophysiology^23^, and pharmacological agents targeting this system are used to modulate associated symptoms^24^. However, these medications have limited efficacy and tremendous side effects. The development of new medications has been slow, in part, due to the lack of a representative experimental system. Using mouse models carrying a schizophrenia-associated genetic deletion allows us to investigate dopamine dysfunction as a downstream consequence of bona fide genetic risk, directly linking dopaminergic abnormalities to disease etiology without artificially inducing circuit perturbations. The ∼40-fold increased risk of SZ conferred by the 3q29 deletion in humans offers a powerful perturbation to model for investigating the underlying biology of the disorder and identifying novel therapeutic targets^1^. In this study, we show that the 3q29 deletion mouse model has elevated evoked dopamine in the dorsal striatum due to aberrant clearance caused by dopamine transporter dysfunction. The strength of this study is two-fold: we have confirmed that the 3q29 deletion mouse model has face validity making it a powerful tool in not only understanding 3q29 deletion-associated SZ, but also to understand idiopathic SZ. Further, we identified a novel mechanism underlying the hyperdopaminergic state associated with 3q29 deletion. This finding is particularly significant given the high rates of treatment resistance observed in individuals with 3q29 deletion^25–27^, underscoring the need for new therapeutic strategies.

Our findings of selectively elevated dopamine signaling in the dorsal striatum are highly consistent with the clinical literature, lending construct validity to our findings. Neuroimaging studies in patients with SZ have repeatedly demonstrated increased dopamine synthesis capacity and release within associative regions of the dorsal striatum, with these abnormalities strongly linked to psychosis and positive symptom severity^24,28^. The regional specificity observed in our 3q29 deletion model therefore mirrors core dopaminergic alterations identified in patients, supporting the translational relevance of this genetic model for studying schizophrenia-related dopamine dysfunction.

While it has previously been suggested that changes in DAT might be underlying the hyperdopaminergic state in SZ, the clinical literature is mixed. For example, meta-analyses of neuroimaging measuring DAT concluded there were no significant changes in levels of DAT in the striatum of patients with schizophrenia^29,30^. These studies align with postmortem brain studies using tissue homogenates that have reported no changes in DAT in the striatum from patients with schizophrenia^31,32^. Further, genetic alterations of the dopamine transporter itself have not been linked to schizophrenia^33^. However, studies using in situ radioligand binding and autoradiography show significantly lower levels of DAT, selectively in the striatum^13^. Our current findings show that overall protein levels of DAT in the dorsal striatum (as seen in analysis of the DAT and pDAT protein expression) are not changed in the 3q29 deletion model but that membrane-DAT (as determined by radioligand binding) or DAT affinity is decreased, which could help explain the discrepancies in these clinical studies. Another likely cause of discrepancies in human clinical studies is the extreme etiologic heterogeneity in schizophrenia^36,37^. Our focus on a single genetic variant of high effect sharply diminishes this heterogeneity and is a strength of this study.

A point of curiosity in this study is the observed attenuated sensitivity to amphetamine-induced locomotor activity. Chronic amphetamine exposure can induce SZ-like symptoms and may further worsen symptoms in individuals with an existing SZ diagnosis^38^. Further, enhanced amphetamine-induced locomotion is frequently used to validate rodent models of SZ and is often considered a hallmark behavioral phenotype of these models^39,40^. That said, some animal models of schizophrenia exhibit the opposite phenotype, with reduced amphetamine-induced locomotion. One example is the M5 muscarinic receptor mutant mouse, which carries a deletion of the M5 gene located within a genomic region implicated in schizophrenia in humans^41^. Given the known interactions between the M5 muscarinic receptor and DAT^42^, along with the similar alterations in amphetamine-induced locomotor responses, further investigation into the role of M5 in the 3q29 deletion mouse model is warranted.

In the current manuscript, we found changes in PKCβ function in the dorsal striatum of 3q29 deletion mice, which we hypothesize are underlying the downregulation in DAT function and ultimately the hyperpaminergic state. PKCβ has previously been proposed as a therapeutic target in SZ, not for the treatment of core symptoms themselves, but rather as an adjunctive strategy to mitigate antipsychotic-induced weight gain^43^. However, our research suggests that PKCβ may be a viable treatment target for many of the core symptoms of SZ, and as such demands further exploration.

It has been well-documented that untreated schizophrenia results in negative outcomes, and the longer the duration of ineffective treatment, the worse the outcomes can be. Thus, it is critical to rapidly find an effective treatment regimen. Underlying genetic etiology, and associated mechanism of disease, is one major characteristic that may be predictive of response to treatment. In limited cases this has already been proven to be effective^44^. The results presented here, suggesting PKCβ as a treatment target for 3q29 deletion-related schizophrenia, highlights the power of this approach and supports continued efforts to understand the mechanism of rare schizophrenia-related variants.

A major strength of this study is its focus on a single, well-defined rare genetic variant, allowing us to directly link the 3q29 deletion to specific alterations in dopamine signaling and transporter function. However, an important limitation is that these findings were generated in a mouse model and may not fully capture the complexity of human disease. Future studies using human model systems, such as induced pluripotent stem cell-derived neurons, will be important for validating these findings and determining their translational relevance.

## Acknowledgements

This research was supported by the Intramural Research Program of the National Institutes of Health (NIH), Z1ADA000069 to M.M. The contributions of the NIH author(s) are considered Works of the United States Government. The findings and conclusions presented in this paper are those of the author(s) and do not necessarily reflect the views of the NIH or the U.S. Department of Health and Human Services.

