## Supplemental Files for "Schizophrenia-Linked 3q29 Deletion Produces Hyperdopaminergic Signaling in the Dorsal Striatum"

**Supplementary Figures**

**Supplemental Figure 1.**

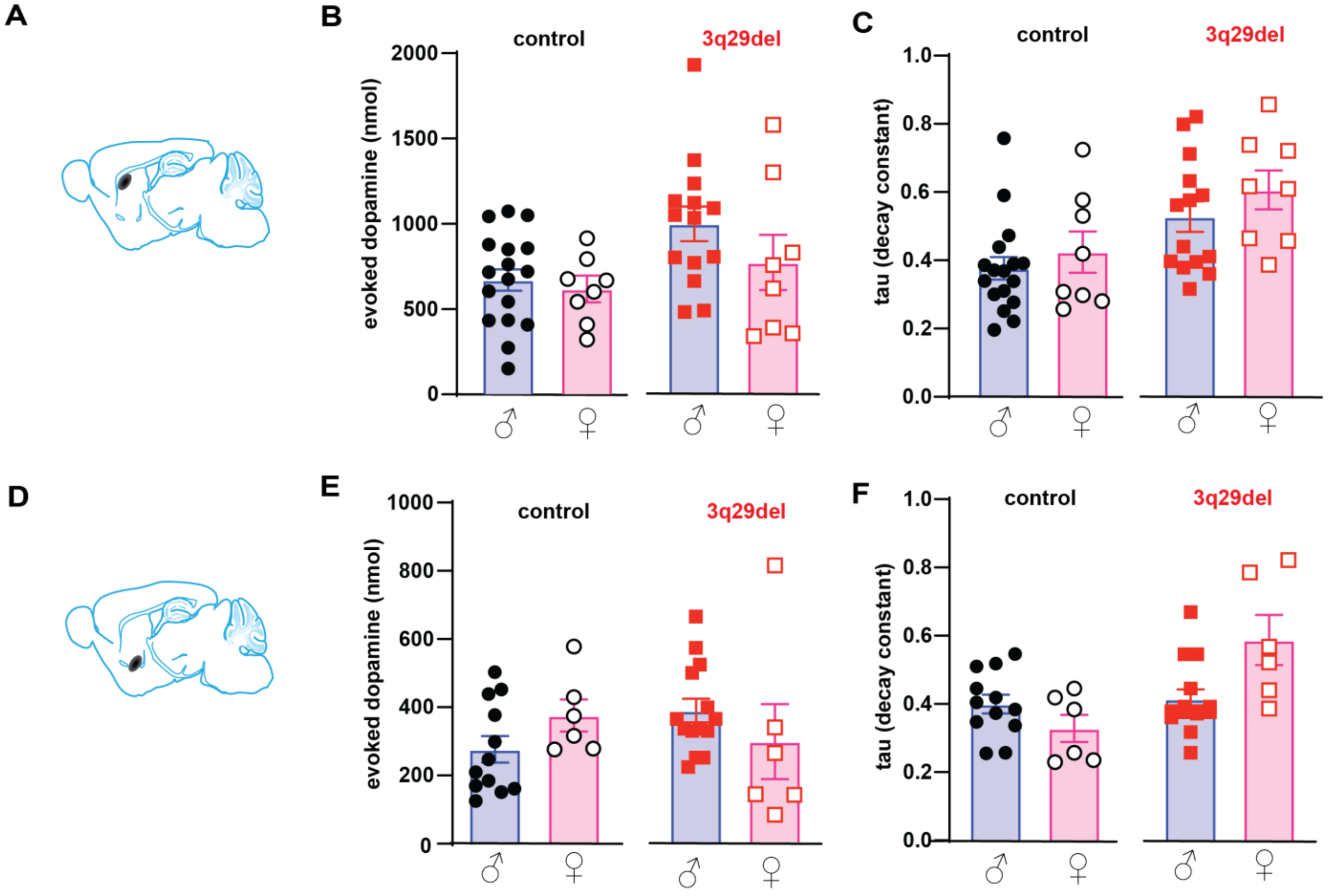


**S1. No sex differences in the dorsal or ventral evoked dopamine measurements across both 3q29deletion mice and control mice. A.** Location diagram of measurement site in dorsal striatum of 3q29del mice (red) compared to control mice (black). **B.** Evoked dopamine in dorsal striatum of male and female control and 3q29 deletion mice. (Two-way ANOVA: main effect of genotype $F(1,43)=4.41$, $p=0.04$, but no main effect of sex $F(1,43)=1.30$, $p=0.26$.) **C.** Decay constant, *tau,* (dopamine uptake efficiency) in the dorsal striatum of male and female 3q29del mice (red) compared to control mice (black). (Two-way ANOVA: main effect of genotype $F(1,43)=12.01$, $p=0.001$, but no main effect of sex $F(1,43)=1.71$, $p=0.20$). **D.** Location diagram of measurement site in nucleus accumbens of 3q29del mice (red) compared to control mice (black). **E.** Evoked dopamine in the nucleus accumbens of male and female control and 3q29 deletion mice. (Two-way ANOVA: no main effect of genotype $F(1,34)=0.05$, $p=0.82$; or sex $F(1,34)=0.005$, $p=0.94$). **F.** Decay constant, *tau,* (dopamine uptake efficiency) in the dorsal striatum of male and female 3q29del mice (red) compared to control mice (black). (Two-way ANOVA: main effect of genotype $F(1,34)=11.51$, $p=0.002$; but not sex $F(1,34)=1.57$, $p=0.22$.) Data is presented as mean ± SEM, with individual points representing data from a single mouse.

### Supplemental Figure 2.

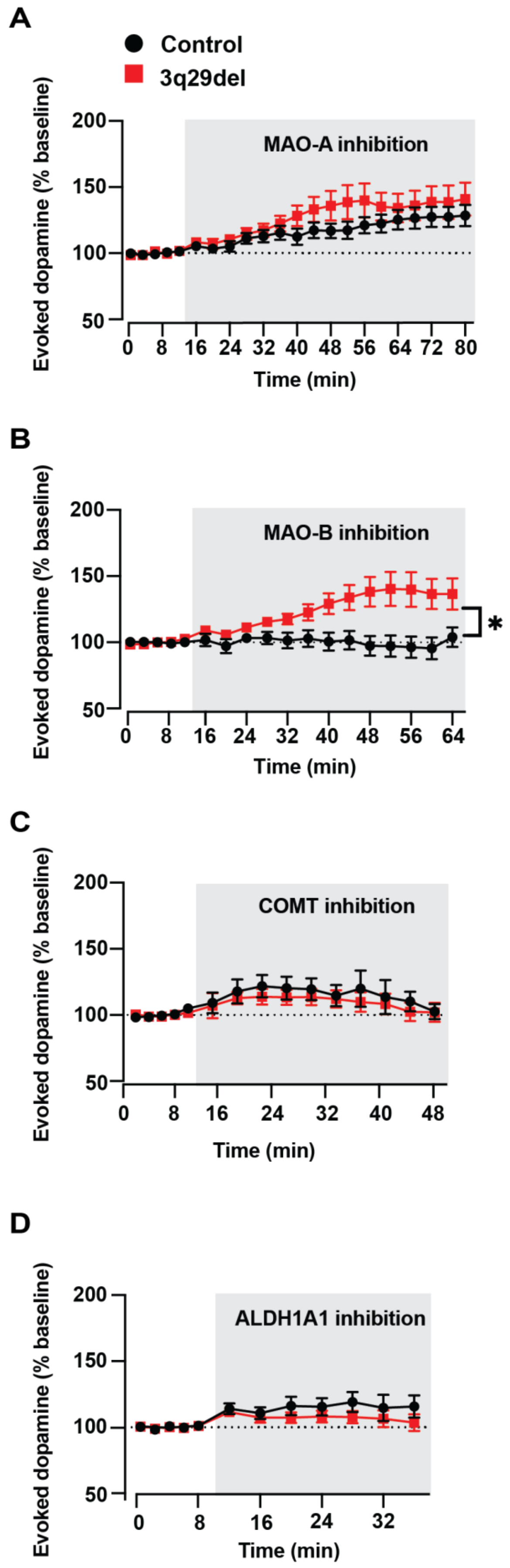


**S2. 3q29 deletion mice show elevated Mao B activity in the dorsal striatum. A.** Time course of dopamine peak amplitudes, normalized to baseline, following application of a Mao-A inhibitor, Clorgiline (100nm), in tissue samples containing the dorsal striatum from 3q29del and control mice (Two-way RM ANOVA: effect of drug application $F(21,679)=15.62$, $p<0.0001$; no effect of genotype $F(1,33)=1.47$, $p=0.23$). **B.** Time course of dopamine peak amplitudes, normalized to baseline, following application of Mao-B inhibitor, Selegiline (100nm), in tissue samples containing the dorsal striatum from 3q29del and control mice (Two-way RM ANOVA: main effect of drug application ($F(17,497)=4.02$, $p<0.0001$; main effect of genotype $F(1,30)=8.82$, $p=0.006$; interaction effect $F(17,497)=5.20$, $p<0.0001$). **C.** Time course of dopamine peak amplitudes, normalized to baseline, following application of a COMT inhibitor, Tolcapone (100nm), in tissue samples containing the dorsal striatum from 3q29del and control mice (Two-way RM ANOVA: main effect of drug application $F(14,370)=3.74$, $p<0.0001$; no main effect of genotype $F(1,28)=0.44$, $p=0.51$). **D.** Time course of dopamine peak amplitudes, normalized to baseline, following application of a ALDH inhibitor, Disulfiram (100µM), in tissue samples containing the dorsal striatum from 3q29del and control mice (Two-way RM ANOVA: main effect of drug application $F(11,441)=5.28$, $p<0.0001$; no main effect of genotype $F(1,41)=1.29$, $p=0.26$). Data is presented as mean ± SEM.

**Supplemental Figure 3.**

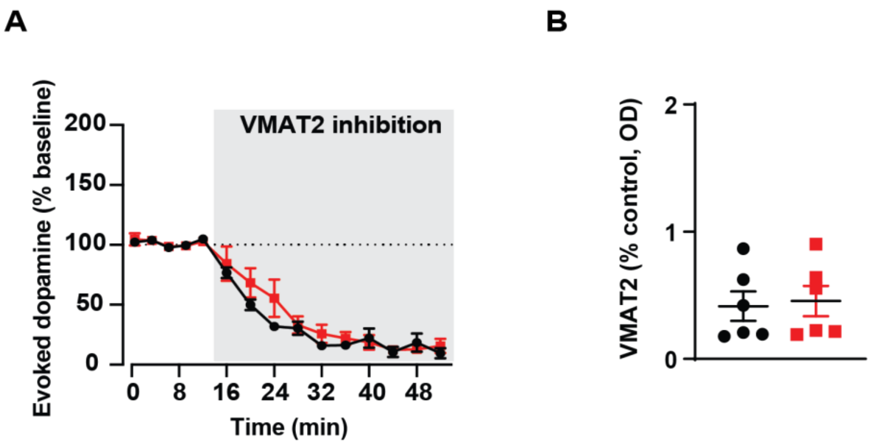


**S3. VMAT function is unaffected in 3q29 deletion mice compared to controls. A.** Time course of dopamine peak amplitudes, normalized to baseline, following application of a VMAT2 inhibitor, tetrabenazine (100µM), in tissue samples containing the dorsal striatum from 3q29del and control mice (Two-way RM ANOVA: main effect of drug application $F(14,163)=76.06$, $p<0.0001$; no effect of genotype $F(1,13)=0.86$, $p=0.37$). **B.** VMAT2 protein, relative to vinculin, in dorsal striatum tissue samples from 3q29del mice (red) compared to control mice (black; $t(10)=0.25$, $p= 0.81$). Data is presented as mean ± SEM, with individual points representing data from a single mouse.

**Supplemental Figure 4.**

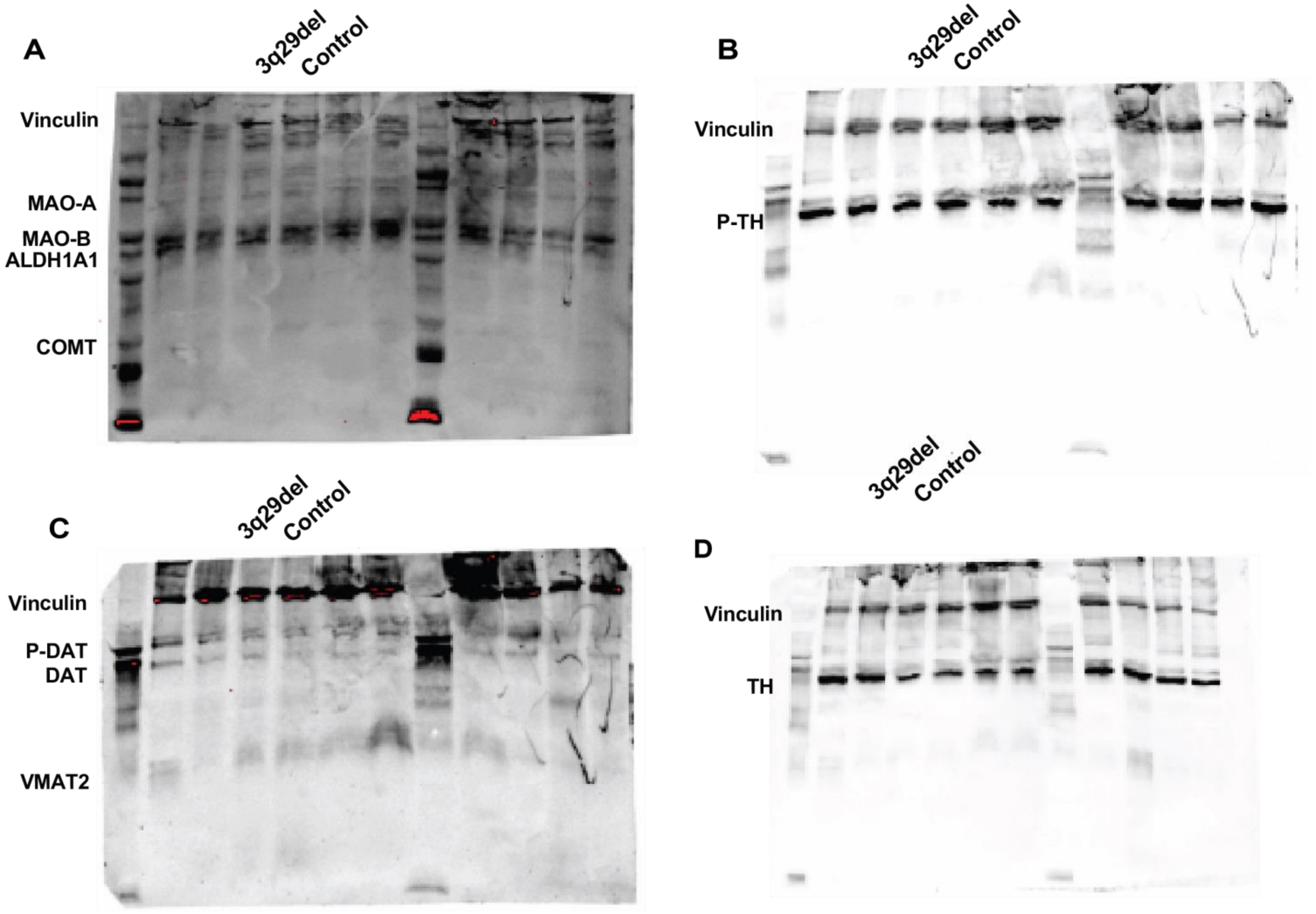


**S4. Representative western blot images.** Samples from the dorsal striatum of control mice compared to 3q29del mice. **A.** Protein bands for vinculin, Mao-A, Mao-B, aldehyde dehydrogenase and COMT are shown. **B.** Protein bands for vinculin and phospho-tyrosine hydroxylase are shown. **C.** Protein bands for vinculin, phosphor- threonine-53- dopamine transporter, dopamine transporter and VMAT2 are shown. **D.** Protein bands of vinculin and tyrosine hydroxylase are shown.

Supplemental Figure 5.

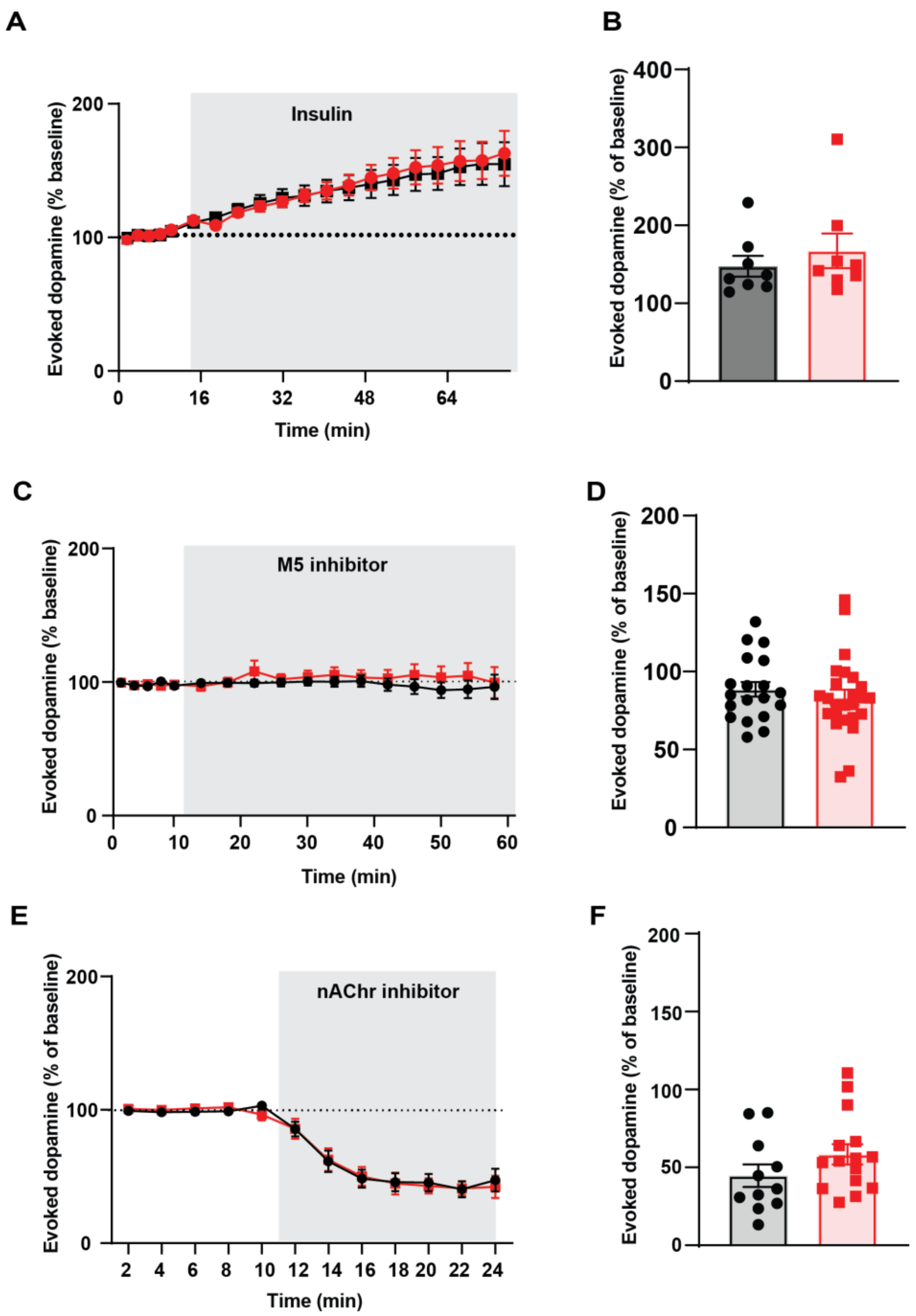


**S5. DAT modulators (insulin, M5, and nAChr) do not lead to differences in dopamine between 3q29 deletion mice and control mice. A.** Time course of dopamine peak amplitudes, normalized to baseline, following application of porcine insulin (50nm; Two-way RM ANOVA: effect of drug application $F(19,261)=24.42$, $p<0.0001$ but not genotype $F(1,14)=0.04$, $p=0.84$). **B.** Following 60 minutes of insulin incubation, no difference was seen in evoked dopamine amplitude between genotypes ($t(14)=0.77$ $p = 0.46$). **C.** Time course of dopamine peak amplitudes, normalized to baseline, following application of the muscarinic receptor 5 antagonist, ML381 (10μM; Two-way RM ANOVA: no effect of drug application $F(16,571)=0.78$, $p=0.71$ or genotype $F(1,41)=0.65$, $p=0.43$). **D.** Following 50 minutes of ML381 incubation, no difference was seen in evoked dopamine amplitude between genotypes ($t(45)=0.39$ $p = 0.70$). **E.** Time course of dopamine peak amplitudes, normalized to baseline, following application of the nAchR inhibitor dihydro-β-erythroidine hydrobromide (DhβE; 1μM). While there was a main effect of the drug application (Two-way RM ANOVA: $F(11,231)=64.12$, $p<0.0001$) there was no effect of genotype ($F(1,22)=0.003$, $p=0.96$). **F.** Following 15 minutes of DhβE incubation, no difference was seen in evoked dopamine amplitude between genotypes ($t(24)=1.4$, $p = 0.17$). Data is presented as mean ± SEM, with individual points representing data from a single mouse.
